# Single-stranded DNA binding protein hitches a ride with the *Escherichia coli* YoaA-χ helicase

**DOI:** 10.1101/2024.06.21.600097

**Authors:** Savannah J. Weeks-Pollenz, Matthew J. Petrides, Robert Davis, Kathryn K. Harris, Linda B. Bloom

## Abstract

The *Escherichia coli* XPD/Rad3-like helicase, YoaA, and DNA polymerase III subunit, χ, are involved in *E. coli* DNA damage tolerance and repair. YoaA and χ promote tolerance to the DNA chain-terminator, 3□-azidothymidine (AZT), and together form the functional helicase complex, YoaA-χ. How YoaA-χ contributes to DNA damage tolerance is not well understood. *E. coli* single-stranded DNA binding protein (SSB) accumulates at stalled replication forks, and the SSB-χ interaction is required to promote AZT tolerance via an unknown mechanism. YoaA-χ and SSB interactions were investigated *in vitro* to better understand this DNA damage tolerance mechanism, and we discovered YoaA-χ and SSB have a functional interaction. SSB confers a substrate-specific effect on the helicase activity of YoaA-χ, barely affecting YoaA-χ on an overhang DNA substrate but inhibiting YoaA-χ on forked DNA. A paralog helicase, DinG, unwinds SSB-bound DNA in a similar manner to YoaA-χ on the substrates tested. Through use of ensemble experiments, we believe SSB binds behind YoaA-χ relative to the DNA ds/ss junction and show via single-molecule assays that SSB translocates along ssDNA with YoaA-χ. This is, to our knowledge, the first demonstration of a mechanoenzyme pulling SSB along ssDNA.

## Introduction

XPD/Rad3-like helicases are critical enzymes for preserving genomic integrity across various organisms (1–5). These superfamily 2 helicases translocate 5□ to 3□ along ssDNA and unwind DNA/DNA and DNA/RNA duplexes in an ATP-dependent manner (2, 6–11). The two *Escherichia coli* XPD/Rad3-like helicases are DinG and YoaA. *In vitro*, YoaA binds χ, a subunit of the DNA polymerase III holoenzyme (pol III HE), to form the active helicase complex YoaA-χ, while DinG functions on its own (11, 12).

The cellular functions of DinG and YoaA-χ are not well understood, and it is therefore, unknown why *E. coli* needs two related helicases. The expression of DinG and YoaA is DNA damage-inducible, indicating DinG and YoaA-χ function in DNA damage repair (13–16). DinG has been implicated in resolving replication and transcription conflicts and unwinds R-loops, D-loops, and G-quadruplexes *in vitro* (9, 17–19). YoaA is less well characterized. YoaA and χ are both required in *E. coli* for promoting tolerance to azidothymidine (AZT), a replication chain terminating agent, and for resisting methyl methanesulfonate (MMS) damage in cells lacking AP endonuclease activity (20, 21).

To understand how YoaA-χ is involved in DNA damage repair, the interactions between YoaA-χ and single-stranded DNA binding protein (SSB) were investigated. SSB binds ssDNA within the cell to protect it from degradation and prevent the formation of secondary structures (reviewed in (22, 23)). SSB also interacts with and coordinates other genomic maintenance proteins, including helicases (reviewed in (23, 24)). YoaA-χ-SSB interactions are of interest because AZT-treated cells form single-stranded (ss) gaps and SSB foci (25). Chi and SSB interactions are also required for χ to promote AZT tolerance, with a mutation at a critical residue on χ for SSB-binding (χ R128A) diminishing AZT tolerance (20). It is unknown if the χ-SSB interaction required for AZT tolerance is when χ is complexed with pol III HE or with YoaA.

To date, interactions between YoaA and SSB have not been identified. However, interactions between SSB and χ are well studied. Chi possesses no catalytic activity and attaches to the pol III HE via the subunit ψ (26, 27). Chi connects the pol III HE to SSB during replication, which promotes stability to the pol III HE (28–30). The last four C-terminal residues of the SSB tail have been crystallized with χψ, showing SSB F177 and χ R128 as critical residues for SSB-χ binding (31). The importance of χ R128 for binding SSB has also been shown via biochemical studies, with the *K_d_* between χ R128A and SSB being too weak to measure compared to wt χψ binding SSB with a *K_d_* of approximately 9 μM (31, 32).

We present here that χ and SSB interact functionally when χ is in complex with YoaA. YoaA-χ-SSB interactions are compared with DinG-SSB interactions due to the previously established interaction between DinG and SSB (33). Both helicases are inhibited by SSB on forked DNA but are indifferent to the presence of SSB on overhang DNA. We also discovered via single-molecule experiments that SSB translocates along ssDNA with YoaA-χ and present evidence of a model where YoaA-χ pulls SSB along DNA like a caboose on a train.

## Results

### SSB has a substrate-specific effect on the DNA unwinding activity of YoaA-χ

To determine if SSB affects the DNA/DNA unwinding of YoaA-χ, the helicase activity of YoaA-χ in the presence and absence of SSB was measured with an *in vitro* FRET-based helicase assay. This assay has been established to be an accurate readout for DNA unwinding activity (11). Briefly, a 20-bp duplexed DNA substrate with a 5□ ssDNA overhang is labeled with the FRET-pair, Cy5 and Cy3, at the blunt end of the duplex. The increase in Cy3 signal with time correlates to the amount of DNA unwound. For these experiments, the DNA substrate contains a 65-nt overhang to allow for both SSB and YoaA-χ to bind. The YoaA-χ ssDNA binding size was estimated using the crystal structure of DinG with a dT_12_ oligo (34). SSB was prebound to DNA before the addition of the helicase. On the 5□ 65-nt overhang substrate (O1), YoaA-χ (50 nM) unwinding activity was barely affected by SSB with a modest stimulation (Fig 1A). A forked DNA substrate containing both 5□ and 3□ overhangs (F1) was also tested. Similar to previously published data, YoaA-χ (50 nM) unwound the forked substrate (F1) faster than the overhang substrate (O1) in the absence of SSB (Fig. 1A vs Fig 1B) (11). Surprisingly, addition of SSB to the forked substrate, which differs from the overhang substrate by the addition of a 3□ 10-nt overhang, inhibited unwinding by YoaA-χ (50 nM) (Fig 1B).

**Fig 1:**
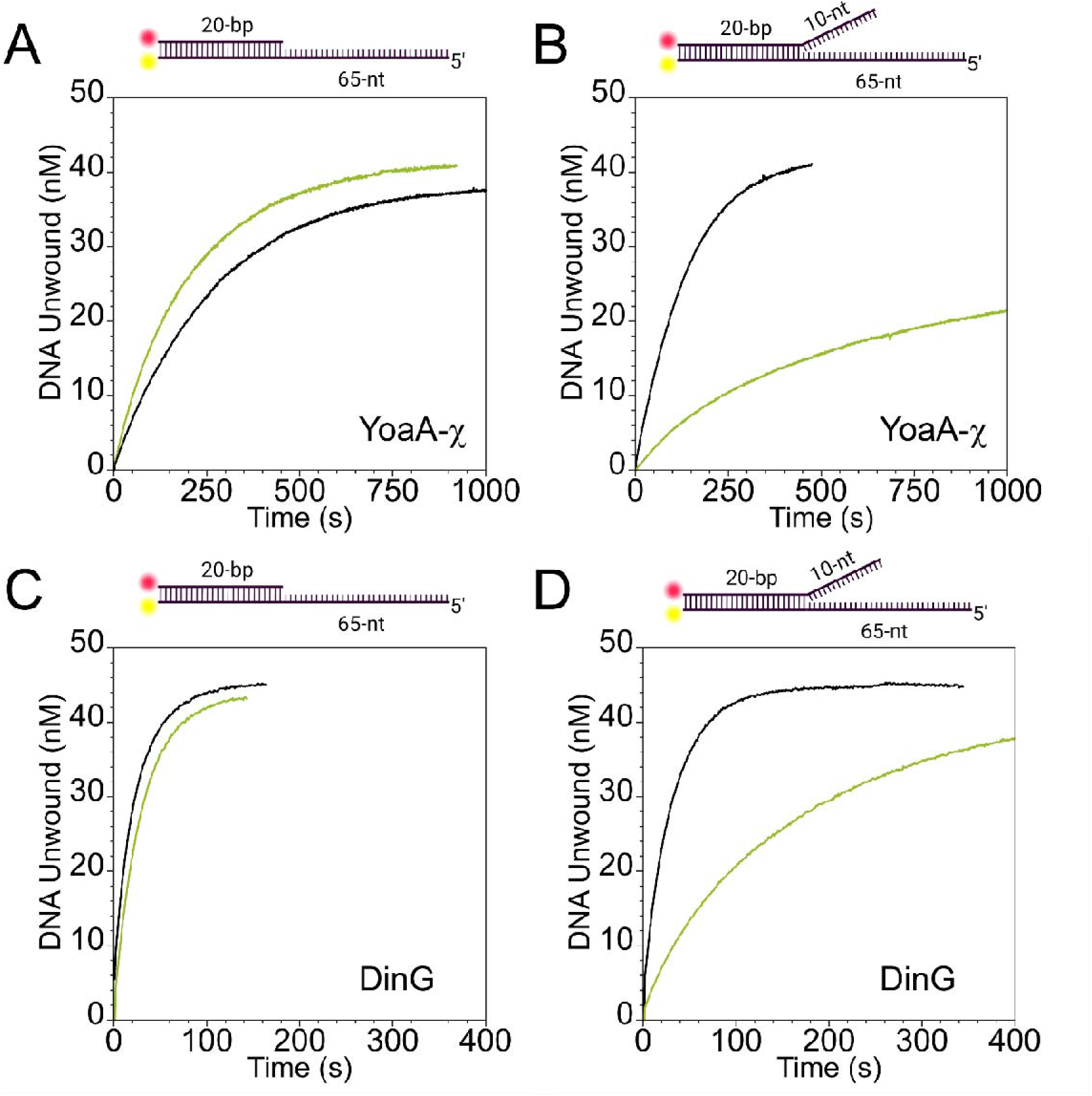
SSB has a substrate-specific effect on helicase activity. Cy5 (red circle) and Cy3 (yellow circle) were attached to the end of a 20-bp duplex DNA containing a 65-nt 5□ overhang with and without a 10-nt 3□ overhang to monitor DNA unwinding. **A.** Time courses of 5□ overhang DNA duplex (50 nM, O1) unwound by YoaA-χ (50 nM) without SSB present (black) and with SSB (75 nM) pre-bound to the DNA (green). **B**. Time courses of forked DNA duplex (50 nM, F1) with a 3□ overhang unwound by YoaA-χ (50 nM) without SSB present (black) and with SSB (75 nM) pre-bound to the DNA (green). **C.** Time courses of 5□ overhang DNA duplex (50 nM, O1) unwound by DinG (5 nM) without SSB present (black) and with SSB (75 nM) pre-bound to the DNA (green). **D**. Time courses of forked DNA duplex (50 nM, F1) with a 3□ overhang unwound by DinG (5 nM) without SSB present (black) and with SSB (75 nM) pre-bound to the DNA (green). With and without SSB conditions were performed side-by-side in the same experiment. Three replicates of unwinding reactions as a function of helicase concentration are shown in Sup Fig 1 and 2.

To determine if YoaA-χ interacts with SSB in a similar manner to its paralog, DinG, these FRET-based helicase experiments were performed with DinG. Because DinG unwinds the two substrates faster than YoaA-χ, the FRET-based helicase experiments were performed with a lower concentration of DinG, 5 nM instead of 50 nM. As for YoaA-χ, the presence of SSB does not significantly alter the unwinding activity of DinG (5 nM) on the overhang substrate (O1) (Fig 1C). On the forked substrate (F1), DinG (5 nM) is also inhibited by the addition of SSB (Fig 1D). DinG is a more robust helicase than YoaA-χ on both substrates tested, with 10-fold less DinG unwinding the substrates faster than YoaA-χ. A range of concentrations were tested for YoaA-χ (50 nM, 20 nM, and 10 nM) and DinG (10 nM, 5 nM, and 2 nM) on the overhang and forked substrates with and without SSB and showed the same phenomena (Sup Fig 1 and 2). SSB alone does not unwind the FRET substrates. The Cy3 signal did not change when only SSB, ATP, and DNA were in the reaction (data not shown).

### The physical interaction between YoaA-**χ** and SSB is necessary for YoaA-**χ** to unwind SSB-bound DNA

To determine whether χ-SSB interactions are necessary for YoaA-χ to unwind DNA bound by SSB, the FRET-based helicase assays were performed on the overhang substrate (O1) with mutant proteins that the weaken the SSB-χ interaction. SSB ΔC1 (F177 deleted) was used to disrupt binding on the SSB side (35). YoaA-χ R128A was used to weaken interactions on the helicase side (31). YoaA-χ R128A alone possessed helicase activity comparable to wild-type (wt) YoaA-χ indicating the mutation in χ did not disrupt the helicase activity of YoaA-χ (Fig 2A). Mutating either protein in the YoaA-χ-SSB complex diminished DNA unwinding, more than the inhibition by wt SSB on the forked DNA (Fig 2B and C vs Fig 1B). Unwinding was only detected for either mutant with the highest YoaA-χ concentration tested (50 nM), albeit very little (Fig 2B and C).

**Fig 2:**
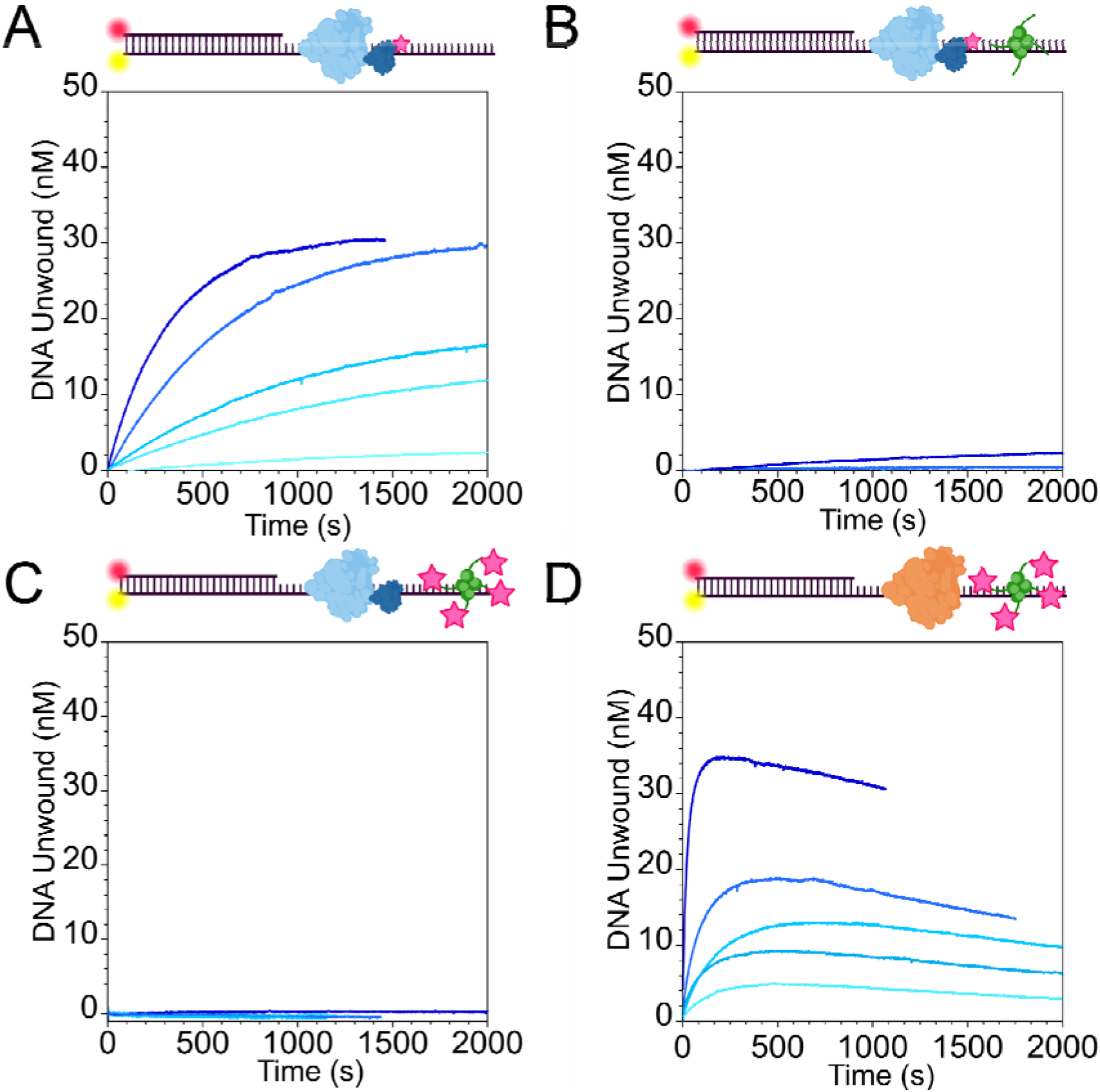
Protein-protein interactions between YoaA-χ and SSB are necessary for YoaA-χ to unwind SSB-bound DNA. Representative time courses for DNA unwinding of 65-nt 5□ overhang DNA (50 nM, O1) are shown for reactions with **A.** no SSB and 2 nM, 5 nM, 10 nM, 20 nM, and 50 nM of YoaA-χ R128A; **B.** wt SSB (75 nM) prebound to DNA and 2 nM, 5 nM, 10 nM, 20 nM, and 50 nM of YoaA-χ R128A; **C.** SSB ΔC1 (75 nM) prebound to DNA and 2 nM, 5 nM, 10 nM, 20 nM, and 50 nM of YoaA-χ. **D.** SSB ΔC1 (75 nM) prebound to DNA and by 2 nM, 5 nM, 10 nM, 20 nM, and 50 nM of DinG. All experiments were performed in triplicate. Darkness in blue corresponds to increasing concentrations of helicase. On the DNA schematic, the red circle indicates Cy5, the yellow circle indicates Cy3, YoaA-χ is denoted in blue, SSB in green, DinG in orange, and pink stars represent protein mutations.

Because it is unknown where SSB binds DinG, only DNA unwinding for wt DinG was measured in the presence of SSB ΔC1 (Fig 2D). DinG unwound more DNA in the presence of SSB ΔC1 than YoaA-χ, though more slowly than when wt SSB was present (Fig 2D vs Fig 1C). At higher concentrations of DinG with SSB ΔC1, Cy3 fluorescence increased due to unwinding followed by a slower decrease, suggesting that DNA was being slowly reannealed (Fig 2D). Using native gel electrophoresis, it was confirmed the overhang substrate was slowly reannealed in reactions with DinG and SSB ΔC1 (Sup Fig 3).

### SSB and YoaA-**χ** bind DNA concurrently

Chi-SSB interactions facilitated unwinding of dsDNA by YoaA-χ when the ssDNA overhang was bound by SSB. To determine whether this is because protein-protein interactions allow both the helicase and SSB to bind DNA at the same time, or because χ-SSB interactions enable YoaA-χ to displace SSB from DNA, electrophoretic mobility shift assays (EMSAs) were performed. For EMSAs, a Cy5-labeled dT_65_ oligo (S11) was used to match the 5□ ssDNA overhang in the FRET substrates and ATPγS to prevent translocation of the helicases. When YoaA-χ (50 nM to 400 nM) was titrated onto the naked dT_65_ oligonucleotide (50 nM), multiple helicase-DNA species were present with as many as 3 to 4 helicases binding dT_65_ (Fig 3A).

**Fig 3:**
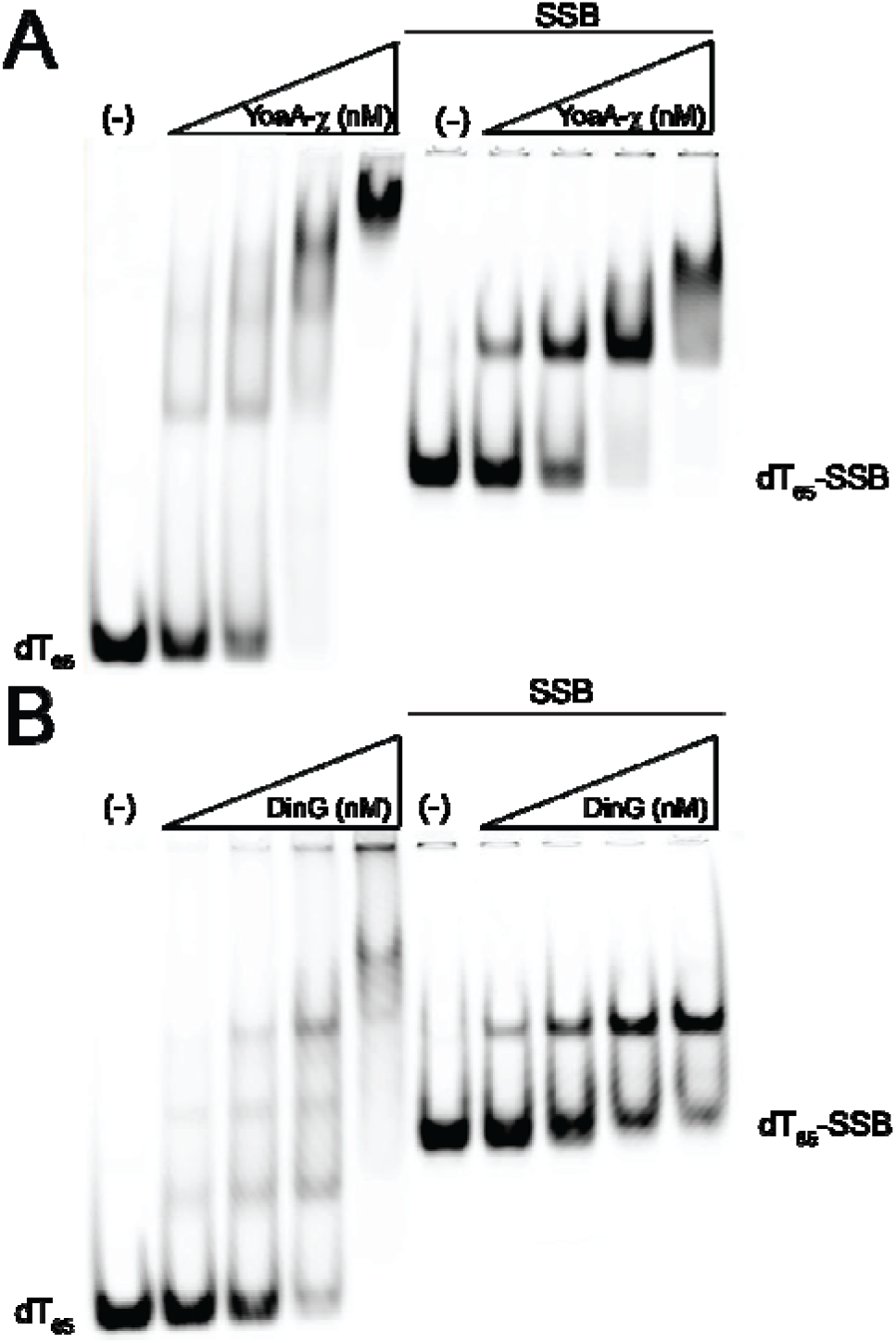
SSB and YoaA-χ bind DNA concurrently. **A.** EMSA of Cy5-labeled dT_65_ oligo (50 nM, S11) titrated with YoaA-χ (50 nM, 100 nM, 200 nM, and 400 nM) with (right panel) and without (left panel) SSB (75 nM) pre-bound to the DNA. **B.** EMSA of Cy5-labeled dT_65_ oligo (50 nM, S11) titrated with DinG (50 nM, 100 nM, 200 nM, and 400 nM) with (right panel) and without (left panel) SSB (75 nM) pre-bound to the DNA. The (-) lanes contain zero helicase in the reaction. The gels shown are representative of three experiments.

When the dT_65_ substrate (S11) was prebound with SSB (75 nM), a second band in addition to the DNA-SSB band was present which likely represents binding of one YoaA-χ along with SSB (Fig 3A). At the highest concentration of YoaA-χ (400 nM) a third band was present, which could be two helicases bound to the DNA-SSB complex (Fig 3A). Similar results were measured for DinG. Multiple DinG molecules bound naked dT_65_ in the absence of SSB and one DinG molecule likely bound in the presence of SSB (Fig 3B). However, YoaA-χ appears to bind SSB-bound dT_65_ with higher affinity than DinG as reflected in the band intensities as a function of helicase concentration. (Fig 3A and B).

### SSB-**χ** interactions are required for YoaA-**χ** to bind DNA-SSB

When χ-SSB interactions were weakened by mutation, YoaA-χ unwinding activity was inhibited on SSB-bound substrates. To determine whether the mutations hinder simultaneous binding of the helicase and SSB to DNA, EMSAs were performed with SSB ΔC1 and χ R128A. When DNA was prebound with SSB ΔC1, binding of YoaA-χ to the dT_65_ oligonucleotide was impaired (Fig 4A). The loss of the SSB ΔC1-only band and smearing in the lanes at the highest two helicase concentrations suggested YoaA-χ bound weakly under these conditions (Fig 4A). When YoaA-χ R128A was tested with wt SSB, YoaA-χ R128A bound the DNA-SSB complex at the highest concentrations, but to a lesser degree compared to wt YoaA-χ (Fig 4B vs Fig 3A). YoaA-χ R128A bound naked dT_65_ DNA similarly to wt YoaA-χ, showing that the mutation in χ did not affect the DNA binding activity of the YoaA-χ helicase (Fig 4A and B left panels). SSB ΔC1-bound to dT_65_ DNA also inhibited DinG binding (Fig 4C). It is interesting to note that two bands were present in the no helicase (-) lanes that only contained SSB ΔC1 and dT_65_ DNA, indicating that deleting F177 altered SSB ΔC1-DNA binding compared to wt SSB-DNA (Fig 4A and C (-) lanes).

**Fig 4:**
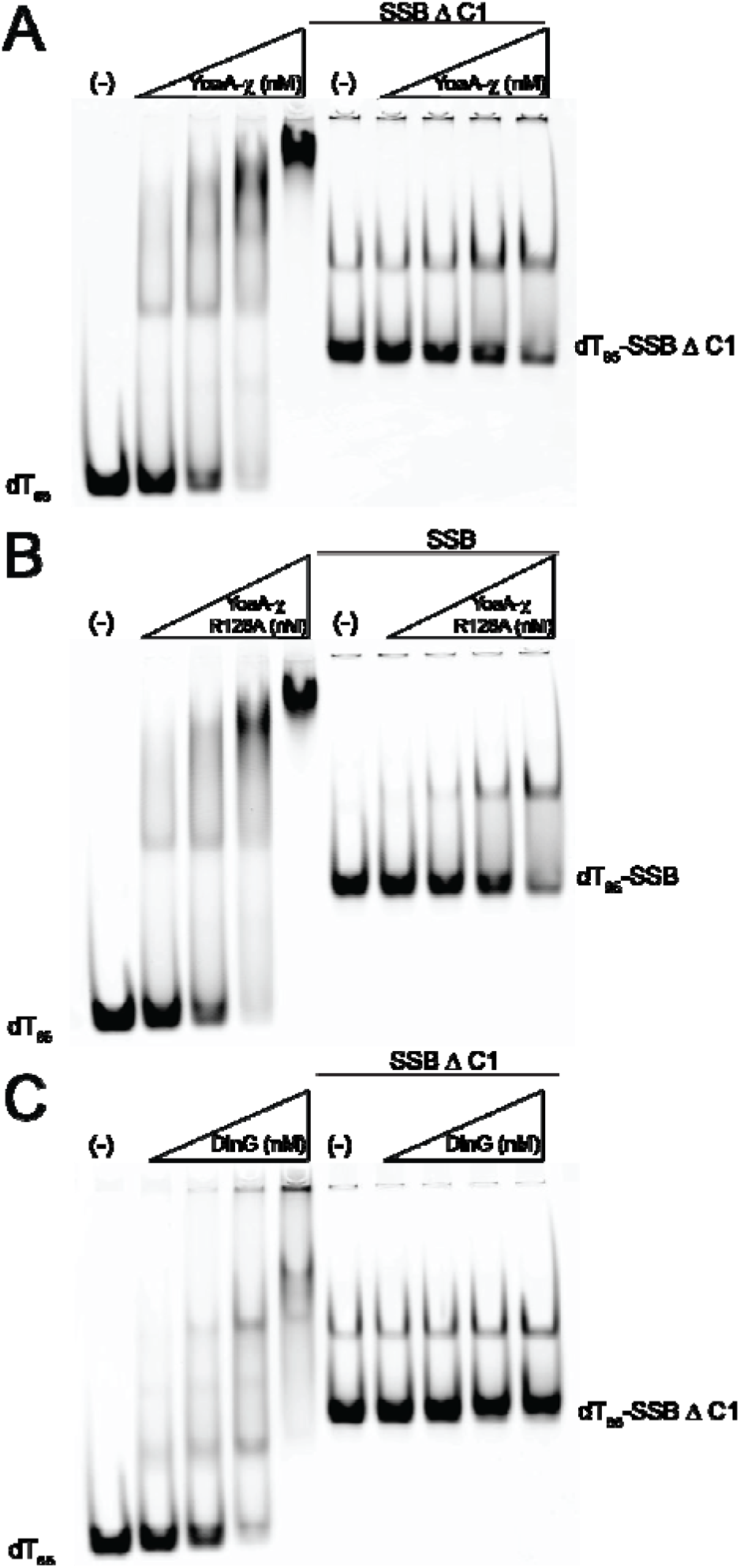
SSB-χ interactions are required for YoaA-χ to bind DNA-SSB. Proteins binding to Cy5-labeled dT_65_ oligo (50 nM, S11) were measured in assays with **A.** YoaA-χ (50 nM, 100 nM, 200 nM, and 400 nM) with (right panel) and without (left panel) SSB ΔC1 (75 nM) pre-bound to the DNA; **B.** YoaA-χ R128A (50 nM, 100 nM, 200 nM, and 400 nM) with (right panel) and without (left panel) wt SSB (75 nM) pre-bound to the DNA; **C.** DinG (50 nM, 100 nM, 200 nM, and 400 nM) with (right panel) and without (left panel) SSB ΔC1 (75 nM) pre-bound to the DNA. The (-) lanes contain zero helicase in the reaction. The gels shown are representative of three experiments.

### Position of YoaA-**χ** in relation to SSB on DNA

To better understand YoaA-χ and SSB interactions, the arrangement of YoaA-χ and SSB on DNA was investigated. The position of YoaA-χ on DNA was measured using the distant-dependent quench that iron-sulfur (Fe-S) clusters in XPD/Rad3 helicases have on fluorophores (11, 36–38). YoaA-χ quenches fluorescein in a distance-dependent manner when YoaA-χ is bound to a 5□ 35-nt overhang and fluorescein is located on a 30-bp duplex region (11). To allow room for both SSB and helicase to bind, a DNA substrate with a longer 65-nt 5□ ss overhang (O3) was used. YoaA-χ (1 μM) and DinG (1 μm) alone quenched fluorescein in a distant-dependent manner on this DNA when fluorescein was in the duplex region either 4-, 7-, 11-, 16-, or 20-nt from the ds/ss junction (Fig 5A). ATPγS was used in the fluorescein quench assays to prevent translocation and unwinding.

**Fig 5:**
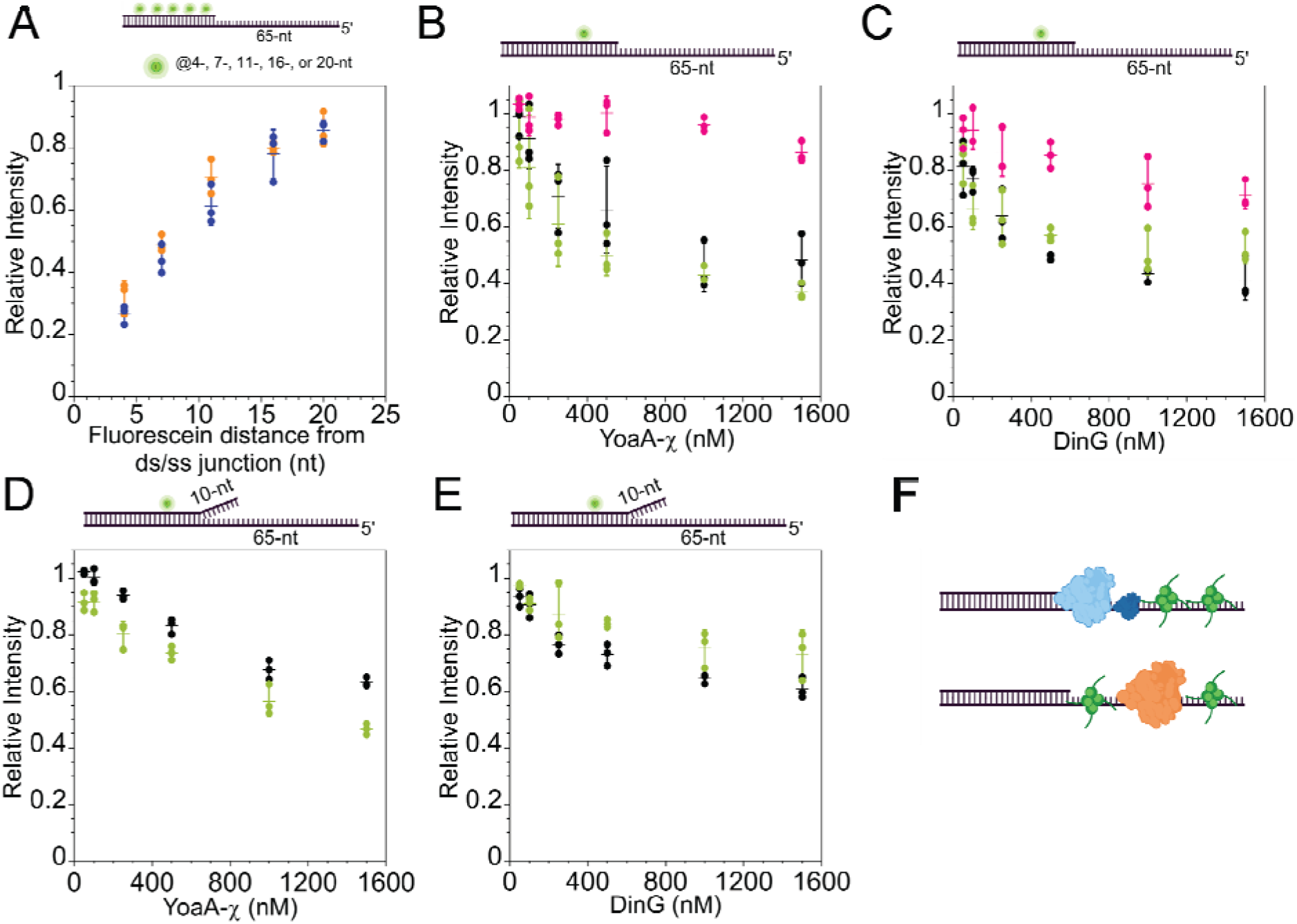
Relative positions of the helicases and SSB on DNA. **A.** Relative intensity of fluorescein (glowing green circle) located either 4-, 7-, 11-, 16-, or 20-nt away from ds/ss junction on an overhang substrate (50 nM, O3) in the presence of either 1 µM YoaA-χ (blue) or DinG (orange). Relative intensity of fluorescein located 7-nt away from the ds/ss junction on an overhang substrate (50 nM, O4) is shown when **B.** YoaA-χ (50 nM, 100 nM, 250 nM, 500 nM, 1 μM, or 1.5 μM or **C.** DinG (50 nM, 100 nM, 250 nM, 500 nM, 1 μM, or 1.5 μM) is added. Relative intensity of fluorescein located 7-nt away from the ds/ss junction on a forked substrate (50 nM, F2) is shown when **D.** YoaA-χ (50 nM, 100 nM, 250 nM, 500 nM, 1 μM, or 1.5 μM) or **E.** DinG (50 nM, 100 nM, 250 nM, 500 nM, 1 μM, or 1.5 μM) is added. For panels **B** – **E**, black indicates naked DNA, green indicates SSB (75 nM) prebound to DNA, and pink indicates SSB ΔC1 (75 nM) prebound to DNA. Dots represent individual experiments, dashed lines represent averages of these three experiments, and error bars indicate standard deviation. **F.** Model of the position of SSB (green) on DNA in relation to YoaA-χ (light and dark blue) and DinG (orange).

To determine the position of YoaA-χ in relation to SSB, the DNA substrate with fluorescein located 7-nt away from the ds/ss junction (O4) and 65-nt ss overhang was utilized. SSB itself quenches fluorescein by directly interacting with the fluorophore (39). When fluorescein is 7-nt away from the ds/ss junction fluorescence is affected by the presence of YoaA-χ but not SSB (Sup Fig 4A). The rationale for these experiments is that if SSB is located between YoaA-χ and the ds/ss junction, YoaA-χ would exert a smaller quench on fluorescein because SSB is increasing the average distance of YoaA-χ from the fluorophore. If YoaA-χ is located between SSB and the ds/ss junction, a larger quench on fluorescein caused by the helicase being bound on average closer to the fluorophore is expected. Because the level of fluorescein quenching is dependent on both the fraction of DNA molecules bound by the helicase and distance of the helicase Fe-S cluster from fluorescein, YoaA-χ (50 nM – 1.5 μM) was titrated on the DNA substrate (50 nM, O4) in the absence and presence of SSB (75 nM) (Fig 5B black without SSB and green with SSB, Table S4). If SSB was present, it was preincubated with DNA. In the absence of SSB, YoaA-χ quenched the intensity of fluorescein in a concentration-dependent manner, with higher concentrations of YoaA-χ correlating with smaller fluorescein intensities (Fig 5B, black, Table S4). In the presence of SSB, YoaA-χ had a slightly larger quench on fluorescein at all the concentrations tested (Fig 5B, green, Table S4). With and without SSB, the relative fluorescein intensity as a function of YoaA-χ concentration produced a hyperbolic-shaped curve that leveled off around 1 μM YoaA-χ (Fig 5B, black without SSB and green with SSB).

DinG had a similar effect on the intensity of fluorescein as YoaA-χ in the absence of SSB (Fig 5C, black, Table S4). Increasing concentrations of DinG increased the quench on fluorescein, with the level of fluorescein intensity leveling off around 1 μM DinG (Fig 5C, black). In the presence of SSB, DinG still quenched fluorescein in a concentration dependent manner (Fig 5C, green). However, in contrast to YoaA-χ, the presence of SSB resulted in a smaller quench on fluorescein by DinG at the higher concentrations of DinG tested (Fig 5C, green, Table S4). Albeit the differences in fluorescein intensity with and without SSB present were small for both helicases, SSB consistently increased the quench by YoaA-χ and decreased the quench by DinG (Fig 5B and C). We believe this is evidence YoaA-χ and DinG arrange with SSB along DNA in opposite ways and may indicate SSB binds behind YoaA-χ and in front of DinG (Fig 5F).

The arrangement of these helicases was also measured in the presence of SSB ΔC1. It was confirmed that SSB ΔC1 alone does not affect the fluorescence of fluorescein located 7-nt away from the junction (Sup Fig 4A). In the presence of SSB ΔC1, both helicases had a smaller quench on fluorescein (Fig 5B and C, pink, Table S4).

### YoaA-**χ** and SSB positions are maintained on forked DNA

One explanation for why both helicases are inhibited on the forked DNA by SSB could be due to SSB binding both ssDNA arms of the fork at the same time. To test this model, the fluorescein quench method was used to determine if SSB was binding at the fork between the helicases and the fluorescein. The fluorescein-labeled forked substrate (F2) has the same sequence for the 3□ overhang as the 3□ overhang on the FRET substrate to negate sequence-dependent effects. We again confirmed wt SSB did not quench the fluorescein on the forked substrate (F2, Sup Fig 4B). Titrating YoaA-χ without SSB on the forked substrate (F2) generated a concentration-dependent quench on fluorescein, but relative intensity of fluorescein leveled off at a higher value, though, than for YoaA-χ on the overhang substrate (Fig 5D, black vs B, black). This suggests that YoaA-χ is binding on average farther from fluorescein on the forked substrate than the overhang substrate. YoaA-χ caused a larger quench on fluorescein on the forked substrate when SSB was bound, similar to the effect of SSB on the overhang substrate (Fig 5D green). Therefore, SSB is not likely being sequestered at the ds/ss junction on the forked substrate and blocking YoaA-χ from accessing the junction. Interestingly, in the presence of SSB, the relationship between YoaA-χ concentration and fluorescein intensity appeared more linear than hyperbolic, indicating the presence of SSB on the forked substrate increased the saturation point of YoaA-χ (Fig 5D, green).

As for DinG, in the absence of SSB, DinG produced a concentration-dependent quench on fluorescein that was weaker compared to the overhang substrate (Fig 5E, black vs C, black). In the presence of SSB, DinG quenched the fluorophore even less (Fig 5E, green). DinG, with and without SSB, still produced a hyperbolic curve. (Fig 5E). This is further evidence that the helicases arrange with SSB differently along the DNA, as illustrated in Figure 5F, and their arrangement with SSB does not differ between an overhang and forked substrate. Both helicases exerted a smaller quench on fluorescein on the forked substrate versus the overhang substrate in the absence of SSB (Fig 5). Perhaps, the 3□ overhang interferes with proteins binding near the ds/ss junction such that the helicases are on average farther from fluorescein and the effective length of the 5□ overhang is shorter than 65-nt. Alternatively, the 10-nt 3□ overhang may provide an extra binding site for the helicases, placing the Fe-S cluster farther away from the fluorophore.

### YoaA-**χ** pulls SSB while it translocates

None of the experiments performed so far reveals the fate of SSB during YoaA-χ translocation. EMSAs showed SSB is still bound to DNA when YoaA-χ binds, but once YoaA-χ translocates, it is unknown if SSB remains or is pushed off the ssDNA. In single-molecule experiments, Alexa Fluor 647-labeled SSB was visualized on ssDNA using confocal fluorescence in the absence and presence of unlabeled YoaA-χ. These experiments were performed by tethering biotinylated ssDNA (20, 452-nt) between two streptavidin-coated polystyrene beads (Fig 6). An experimentally determined concentration of SSB-AF647 (20 pM) that does not saturate the DNA was added (Fig 6A). As expected, SSB-AF647 in our assay conditions bound to the ssDNA while held at a constant force of 10 pN (Fig 6A) (40, 41). When SSB alone bound ssDNA (number of binding events (n) = 442 on 22 DNA molecules), short binding events, most of which had a lifetime of 2 s or less, were measured (Fig. 6D). The longest binding events lasted for 20 seconds. To accurately obtain rates of SSB movement, only tracks longer than 4 seconds were included in calculations to minimize noise from shorter tracks. Only 8 tracks were longer than 4 seconds, with an average slope of 0.010 ± 0.004 µm/sec. These tracks appeared to exhibit inconsistent, diffusive motion and did not move systematically in any direction. This agrees with previous literature, where it is well-documented SSB freely diffuses on ssDNA in single-molecule experiments (42).

**Fig 6:**
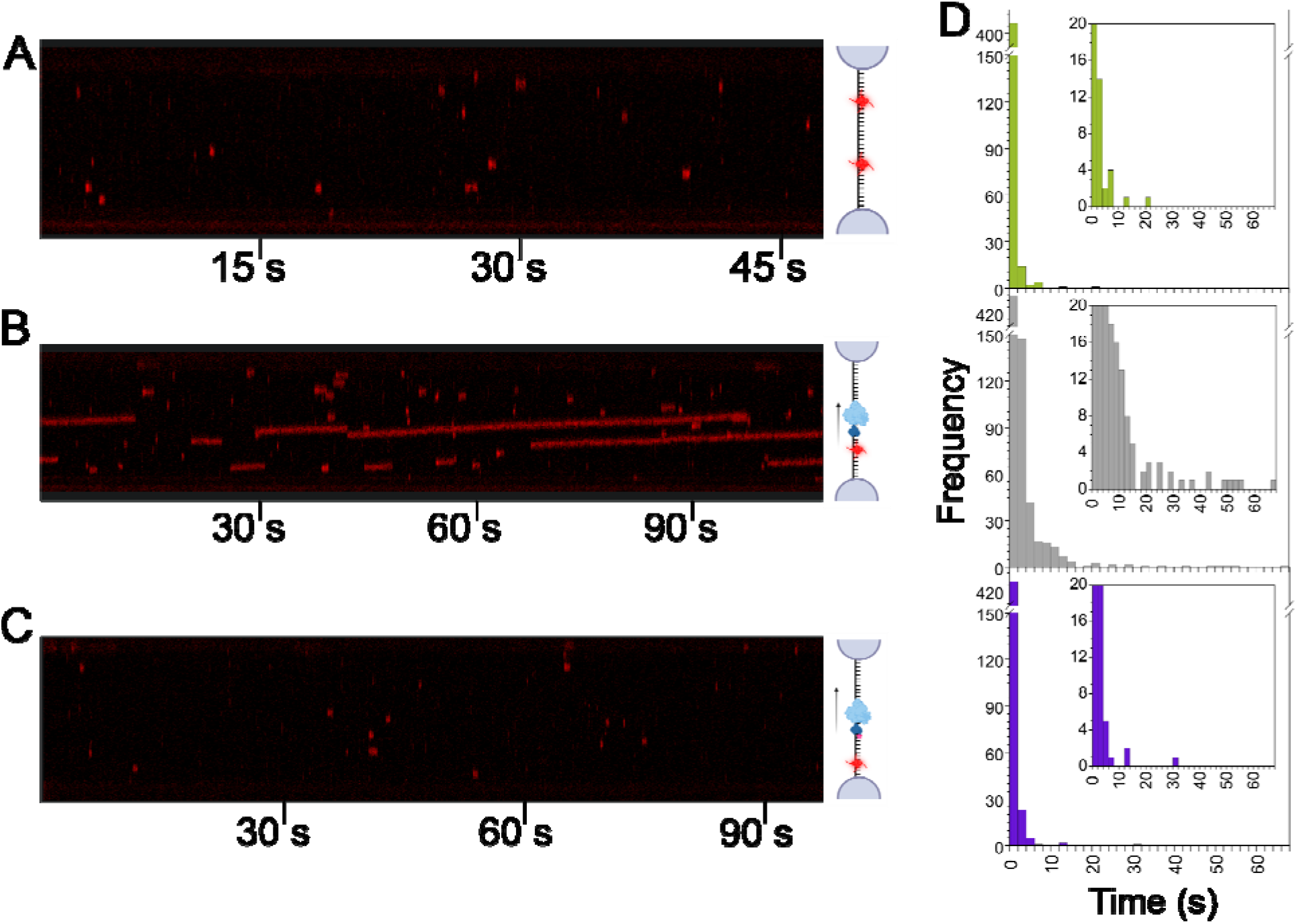
YoaA-χ pulls SSB while it translocates along ssDNA. All kymographs were collected with biotinylated ssDNA (20,452-nt) tethered between two streptavidin beads at constant force of 10 pN. **A.** Representative kymograph of ssDNA in a channel containing AF647-labeled SSB (red, 20 pM) and ATP (4 mM). **B.** A kymograph is shown for the same DNA strand that was used in panel **A.** and moved to a different channel containing AF647-SSB (red, 20 pM), YoaA-χ (100 nM), and ATP (4 mM). **C.** Representative kymograph of ssDNA in a channel with AF647-labeled SSB (red, 20 pM), ATP (4 mM), and YoaA-χ R128A (100 nM). **D.** Binding events of each experimental condition were binned into intervals of 2 seconds and shown in histograms. The SSB negative control (top) contained 442 total tracks measured on 22 different DNA molecules. Wild-type YoaA-χ (middle) contained 745 total tracks measured on 15 different DNA molecules. YoaA-χ R128A (bottom) contained 490 total tracks measured on 35 different DNA molecules.

To determine if YoaA-χ removes SSB from DNA when it translocates, 100 nM YoaA-χ was added to 20-knt ssDNA with 20 pM SSB-AF647 (Fig 6B). In the presence of YoaA-χ, SSB binding events increased in frequency (n = 745 on 15 DNA molecules) (Fig 6D). The SSB was not stripped from the DNA and, surprisingly, was even retained on the DNA for longer time periods, as shown by longer lifetime binding events. With wt YoaA-χ, SSB lifetimes on DNA were as long as 67 seconds (Fig 6D). Unlike in the case of SSB binding DNA alone, many longer SSB tracks in the presence of YoaA-χ were visibly sloped over their duration. There were 115 tracks longer than 4 seconds, with an average slope of 0.023 ± 0.001 µm/sec. Critically, all these tracks sloped in the same direction within each kymograph, which matches what is expected of the 5□ to 3□ unidirectional movement of YoaA-χ. This suggests that SSB does not fall off the ssDNA while YoaA-χ translocates, but instead actually moves with the helicase.

To determine if the movement of SSB with YoaA-χ is due to the physical interaction between χ and SSB, the same experiment was performed with YoaA-χ R128A (Fig 6C). With YoaA-χ R128A, SSB exhibited shorter lifetimes on DNA and only a few SSB molecules moved with the helicase (Fig 6D). This condition had the least number of SSB binding events per DNA molecule, with n= 490 on 35 DNA molecules, and kymographs were similar to SSB only kymographs (Fig 6D). The longest track measured was 31 seconds, with only 9 tracks longer than 4 seconds and an average slope of 0.024 ± 0.004 µm/sec (Fig 6D). These longer tracks and their slopes closely resembled those of SSB with wt YoaA-χ and is likely due to YoaA-χ R128A movement. These data indicate that the physical interaction between χ and SSB is required for SSB to move along the DNA with YoaA-χ.

## Discussion

SSB caused a substrate-specific effect on the DNA/DNA unwinding activity of YoaA-χ and the YoaA paralog, DinG. The unwinding activity of both helicases was not significantly affected by the presence of SSB on an overhang substrate, although there was minimal stimulation of YoaA-χ. This slight SSB-induced stimulation of YoaA-χ on overhang DNA is significant because SSB occludes a portion of the DNA YoaA-χ binds and could block access to the ssDNA or ds/ss junction, yet YoaA-χ is still stimulated despite this. In contrast to the overhang substrate, both helicases were inhibited on forked DNA, which is intriguing since any overhang structure becomes forked as it is unwound. It is possible our overhang substrate may possess a duplex length (20-bp) insufficient for SSB to become inhibitory during unwinding. Physiologically, this SSB-induced inhibition on forked DNA may prevent YoaA-χ and DinG from unwinding long stretches of DNA within the cell. If YoaA-χ does act on DNA during AZT damage repair, unwinding long stretches of DNA may be deleterious in this replication-coupled repair system.

Our results regarding SSB’s effect on DinG helicase activity differ from those of Cheng *et al*, in which SSB stimulated the helicase activity of DinG (33). The substrates used may cause this difference in SSB’s effect on DinG, as Cheng *et al*. used a M13mp18 circular ssDNA with a 55-nt primer annealed (33). Perhaps the large ssDNA region of the M13mp18 DNA coated by SSB directs DinG to the ds/ss junction, but more studies must be conducted to resolve these differences.

The relative indifference of YoaA-χ to the presence of SSB on overhang DNA during unwinding is partly caused by the physical interaction between χ and SSB. SSB-χ interactions are well established, with χ R128 being a critical residue for binding SSB (31, 32). We confirmed χ R128A did not disrupt the DNA binding and unwinding activity of the YoaA-χ complex in the absence of SSB. In the presence of SSB, though, YoaA-χ R128A bound ssDNA with lower affinity and barely unwound DNA, indicating the χ-SSB interaction is required for YoaA-χ to bind and unwind SSB-bound DNA. This is to be expected, since physical interactions between SSB and SSB interacting proteins (SIPs) play an important role in eliciting productive SSB-induced effects on SIPs (reviewed in (24)). We propose the physical χ-SSB interaction allows YoaA-χ to rearrange SSB along the DNA to create a DNA binding site for the helicase.

The *E. coli* helicases RecQ and PriA bind SSB-DNA by promoting the SSB 35-nt binding mode over the 65-nt binding mode via direct helicase-SSB interactions (43, 44). Because the 35-nt binding mode of SSB wraps less ssDNA than the 65-nt mode, RecQ and PriA induce the mode switch to expose more ssDNA for the helicases to bind. It is unknown how YoaA-χ rearranges SSB, but YoaA-χ may also be altering the binding mode of SSB or sliding SSB out of the way.

Both YoaA-χ and DinG had weak DNA binding and unwinding activities in the presence of SSB ΔC1. Because SSB ΔC1 is missing the common SIP-interacting residue, F177, which is critical for binding χ and DinG, the helicases’ ability to remodel SSB ΔC1-DNA is likely impaired which leads to inhibition of activity (31, 33). However, the C-terminal tail of SSB also plays a role in SSB-DNA dynamics and we cannot rule out if SSB ΔC1 may inhibit the helicases by also affecting SSB-DNA interactions (44–46). Our EMSAs suggest that SSB ΔC1 binds dT_65_ differently than wt SSB because at equal concentrations two bands were present for SSB ΔC1 compared to one band for wt SSB.

Interestingly, the highest concentrations of YoaA-χ weakly bound SSB ΔC1-DNA in the EMSAs but did not quench fluorescein in the presence of SSB ΔC1. This could be due to SSB ΔC1 binding between YoaA-χ and the ds/ss junction. DinG did not bind SSB ΔC1-DNA in the EMSAs, yet despite this, unwound more DNA in the presence of SSB ΔC1 compared to YoaA-χ. Fluorescein quench results also indicated high concentrations of DinG (≥ 1 μM) quenched fluorescein in the presence of SSB ΔC1 to a higher degree than YoaA-χ. These apparently contradictory results could be explained by the relative strength of each helicase. DinG may be a strong enough motor protein to push SSB ΔC1 to gain access to the ds/ss junction, even though DinG binds SSB ΔC1-DNA weaker than YoaA-χ. This agrees with our FRET-based helicase assays, in which YoaA-χ was a weaker helicase than DinG on the two substrates tested. If DinG pushes SSB ΔC1 along ssDNA via its motor activity, this suggests an active role in remodeling of SSB on DNA.

How SSB inhibits the helicase activity of YoaA-χ and DinG on forked DNA is still unknown. Typically, in the absence of SSB, XPD/Rad3 helicases including YoaA-χ and DinG unwind forked DNA faster than overhang DNA (9, 11, 36, 47, 48). For XPD, this increased activity on forked DNA has been attributed to secondary DNA-binding sites on the helicase (36, 49). YoaA-χ and DinG may also contain a secondary DNA-binding site that binds the displaced strand, and the acidic tip of SSB, which is negatively charged like DNA, could bind this site to inhibit these helicases on forked substrates. This may be a conserved interaction between *E. coli* XPD/Rad3-like helicases that is not shared among the other XPD-like helicases since RPA does not possess the highly conserved acidic tail of *E. coli* SSB. It has been established for *Ferroplasma acidarmanus* (Fac) XPD that a secondary DNA binding site contributes to Fac RPA2’s mechanism of altering XPD helicase activity (50).

Our fluorescein quench data implies SSB binds behind YoaA-χ in relation to the ds/ss junction due to YoaA-χ exerting a stronger quench on fluorescein when SSB was present. The differences in fluorescein intensity on the overhang substrate at the highest concentrations of YoaA-χ, with and without SSB, were small but consistent. Based on the EMSAs, these small differences between the quenching with and without SSB are to be expected. In the absence of SSB, the number of helicase molecules bound to DNA increased to 3 to 4 at high concentrations of YoaA-χ (≥ 200 nM). As the number of helicase molecules on DNA increases, the quench in fluorescence will be larger because on average a helicase molecule is closer to the ds/ss DNA junction. When SSB was bound to the DNA, EMSAs showed only 1 to 2 helicases bound the DNA at high concentrations of YoaA-χ (≥ 200 nM). Therefore, 3 to 4 helicases bound to naked DNA likely quench fluorescein to a similar degree as one helicase and one SSB bound. Our fluorescein quench assays also suggest the opposite arrangement may be true for DinG, with SSB being at the front of the helicase such that DinG could push SSB, due to a weaker quench when SSB is present. Because these are ensemble experiments it is possible these small differences in fluorescein intensity in the presence and absence of SSB reflect a mixture of molecules with SSB binding in front or behind the helicases. Structural data is needed to confirm these predictions of how SSB binds and arranges with respect to both helicases on DNA.

A predicted structure of YoaA-χ bound to a dT_11_ oligo was generated with AlphaFold 3 (51). The predicted YoaA-χ-dT_11_ structure matches the expected orientation of an XDP/Rad3-like helicase on ssDNA. XPD/Rad3-like helicases translocate along DNA with a 5□ to 3□ directionality, which is conferred by the helicase domain I (HDI), arch, and Fe-S domains binding closer to the 3□ end of DNA (36, 49). In accordance with this concept, the top five models generated by AlphaFold 3 for YoaA-χ-dT_11_ all place a dT_11_ oligo within YoaA such that the N-terminal HDI binds the 3□ end of ssDNA, and the C-terminal HDII binds the 5□ end of ssDNA (51). This predicted structure of YoaA-χ-dT_11_ is also similar to the orientation of the solved structure of DinG-dT_12_, with DinG’s HDI binding the 3□ end of DNA and HDII binding closer to the 5□ end (34). The predicted YoaA-χ-dT_11_ structure also places YoaA and χ in an orientation that agrees with published results on YoaA-χ interactions. YoaA R619 and T620, as well as χ F64, are critical residues in YoaA-χ interactions, and the AlphaFold 3 structure situates χ and the HDII of YoaA such that these three residues are within the predicted binding interface (Fig 7) (12, 16).

**Fig 7:**
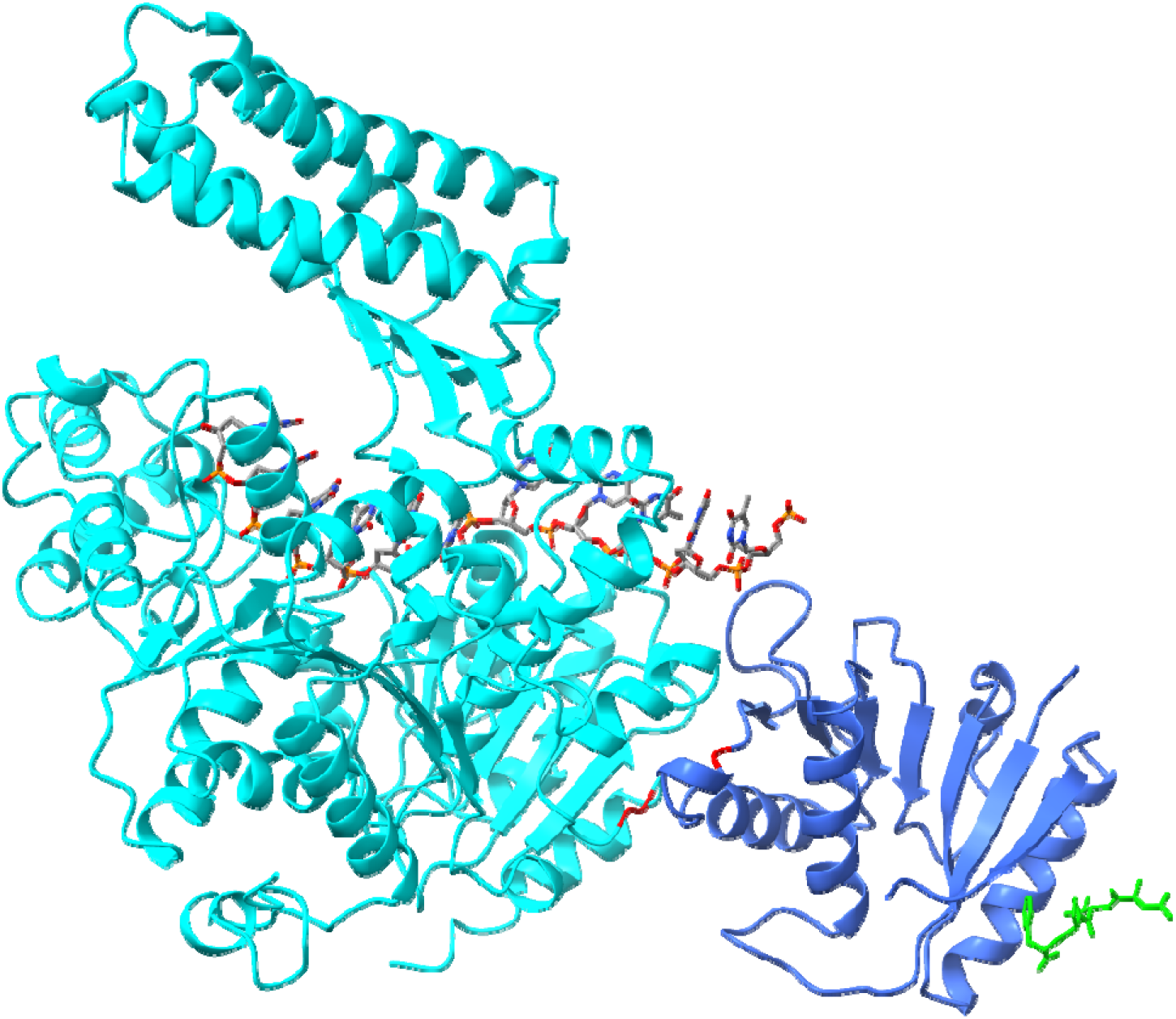
Proposed model of how YoaA-χ binds SSB. AlphaFold 3 prediction of YoaA (light blue) binding a dT_11_ oligo (stick model) and χ (dark blue) (51). AlphaFold 3 predicts χ to bind the C-terminal end of YoaA. The solved crystal structure of χ bound by a C-terminal peptide of SSB (green stick model) is overlayed on this predicted structure (31). Red residues on YoaA indicate R619 and T620, and on χ indicate F64.

To visualize SSB binding on this predicted YoaA-χ-dT_11_ complex, the solved structure of χ bound to an SSB C-terminal peptide was overlaid with the predicted YoaA-χ-dT_11_ structure (31). Because the HDI of XPD/Rad3-like helicases binds closer to the 3□ end of DNA and move 5□ to 3□, the HDI is the “front” of the helicase and encounters the ds/junction first. As shown in Figure 7, SSB is placed near HDII of YoaA and is the “back” of the YoaA-χ complex. Therefore, SSB binding χ makes SSB a “caboose” on the YoaA-χ complex (Fig 7).

Single-molecule results of SSB moving with YoaA-χ also provide evidence towards SSB as the caboose. If SSB bound DNA ahead of YoaA-χ and YoaA-χ pushed SSB, comparable numbers of long sloped tracks with wt YoaA-χ and YoaA-χ R128A are expected. Since there are far fewer long (≥ 4s) tracks with YoaA-χ R128A compared to wt YoaA-χ, and the mutant still possesses translocation activity, this implies SSB binds behind YoaA-χ and is being pulled behind the helicase. The few sloped tracks with YoaA-χ R128A may be due to weak SSB binding to the χ mutant or occasional pushing by helicase.

The single-molecule experiments were performed with the ssDNA held at a constant force of 10 pN, at which SSB binds approximately 17-nt of DNA (40, 41). This weaker wrapping mode of SSB can explain the short-lived and sparse SSB-DNA binding events that occurred with only SSB in the reaction. Motor proteins can also remove SSB off DNA, with the HELQ helicase stripping RPA off ssDNA in single-molecule experiments (52). Due to this weaker SSB binding mode and YoaA-χ being a motor protein, it seems feasible that YoaA-χ could push SSB off the ssDNA. Instead, though, YoaA-χ did not remove SSB from ssDNA during translocation. In fact, along with SSB translocation with YoaA-χ, SSB-DNA binding events per DNA molecule increased in the presence of YoaA-χ, indicating YoaA-χ recruits SSB to ssDNA.

Because we believe YoaA-χ pulls SSB, the possible pushing activity of YoaA-χ does not appear to be the main mechanism for clearing SSB-bound DNA. There are helicases that can push SSB, with Pif1 pushing both *E. coli* SSB and human RPA along ssDNA (53, 54). YoaA-χ may initially access SSB-bound DNA by passive rearrangement between χ and SSB, which is not without precedent. When in complex with the DNA polymerase III clamp loader complex, χ remodels SSB on DNA passively without the use of motor activity (55). However, in the case of YoaA-χ, once the helicase complex gains access to DNA, YoaA-χ actively pulls SSB via its motor activity.

The cellular function of YoaA-χ pulling SSB along on DNA is unknown. FANCJ and RPA coordinate with each other to resolve G-quadruplexes (56–58). Something similar may occur with YoaA-χ and SSB where SSB travels with YoaA-χ to prevent newly unwound secondary-structures from reannealing. SSB may also travel with YoaA-χ to act as a brake once forked DNA is generated. If YoaA-χ is repairing a base-stalling lesion on the leading strand, YoaA-χ would be pulling SSB away from the parental duplex. SSB may need to be moved away from the junction to allow other proteins to bind and access the DNA to repair the lesion. YoaA-χ and SSB may also have a role in fork regression, with SSB being pulled out of the way of other proteins. To our knowledge this is the first report of a mechanoenzyme pulling SSB along DNA and takes the field of how helicases and SSB interact in a new and intriguing direction.

## Materials and Methods

### Buffers

Assay buffer A is 50 mM Tris HCl pH 7.5, 125 mM NaCl, 10 mM MgCl_2_, 50 µg/mL BSA, and 2 mM DTT at final concentration within the reaction. For single-molecule experiments, buffer B is LUMICKS 10x running buffer (PBS, 50 mM sodium azide, and 5 mM EDTA) diluted 1:10 in water and buffer C is buffer B diluted 1:10 in water.

### SDM of *holC* R128A

A Q5 Site-directed Mutagenesis Kit (New England Biolabs) was used per the manufacturer’s instructions with the primers in Table S1 and a pET15b-holC plasmid to mutate arginine 128 to alanine in the HolC protein. DNA sequencing was performed to confirm the desired mutation was made and other mutations were not present.

### Overexpression of proteins

YoaA-χ was overexpressed using a previously published protocol (11). YoaA-χ R128A was expressed in the same manner as the wt helicase. Both preparations of wt and mutant YoaA-χ contain an N-terminal His-tag on YoaA. For DinG, BL21 (DE3) *E. coli* cells were transformed with a modified version of the pET30-his-DinG provided by Dr. Rafael Camerini-Otero in which sequences for the S-tag and factor Xa cleavage site were deleted (8). The modified pET30-his-DinG vector expresses DinG with a N-terminal 6X His tag only. Transformed cells were grown in terrific broth with kanamycin (50 μg/mL) at 37° C shaking at 250 RPM until an OD_600_ of approximately 0.6. To induce protein expression, IPTG (1 mM) was added along with iron supplements, iron (II) sulfate (0.1 mg/mL) and ammonium ferric citrate (0.1 mg/mL). The cells were grown for 4 more hours at 25° C with shaking at 250 RPM. The cells were pelleted by centrifugation at 6,370 RCF for 30 min, media was removed, and the cell pellets stored at −80°C.

The pGN62-SSB (a gift from Dr. Michael O’Donnell) and pET21a-SSBΔC1 (a gift from Dr. James Keck) were transformed into *E. coli* BL21 (DE3) pLysS cells for the overexpression of SSB and SSB ΔC1 respectively (35). SSB ΔC1 denotes the last C-terminal residue, F177, is deleted. The cells were grown in terrific broth with ampicillin (100 μg/mL) and chloramphenicol (30 μg/mL) at 37° C shaking at 250 RPM until an OD_600_ of approximately 0.6. Protein expression was induced with ITPG (1 mM). The cells were grown for 3 more hours at 37° C with shaking at 250 RPM. The cells were pelleted and stored at −80°C.

### Purification of proteins

YoaA-χ was purified by previously published methods and YoaA-χ R128A was purified using the same protocol (11). Two different preparations of purified YoaA-χ R128A were used for all experiments that contained YoaA-χ R128A to confirm the effects of the R128A mutation. For DinG, *E. coli* cell pellets were resuspended in low imidazole buffer (25 mM Tris HCl pH 8, 25 mM imidazole, 1 M NaCl, Sigma FAST Protease Inhibitor Cocktail Tablet EDTA-free, and 0.5 mM TCEP). The cell lysate was loaded onto a HisTrap FF column (Cytiva) and washed with low imidazole buffer. DinG was eluted with a linear gradient from 25 mM to 500 mM imidazole and fractions containing DinG were dialyzed overnight in dialysis buffer A (25 mM Tris HCl pH 8, 1 M NaCl, 0.5 mM TCEP), and then dialyzed for 6 hours in dialysis buffer B (25 mM Tris HCl pH 8, 300 mM NaCl, 0.5 mM TCEP). DinG was loaded onto a HiTrap Heparin HP column (Cytiva). The column was washed with low salt buffer (25 mM Tris HCl pH 8, 300 mM NaCl, and 0.5 mM TCEP) and DinG was eluted with a linear gradient from 300 mM to 1 M NaCl. Fractions containing DinG were dialyzed overnight into storage buffer (25 mM Tris HCl pH 8, 300 mM NaCl, 30% glycerol, and 0.5 mM TCEP) and stored at −80°C. Protein concentration was measured using an A_280_ absorbance of DinG and molar absorptivity of 78,840 M^−1^ cm^−1^. SSB and SSB ΔC1 were purified by previously published methods (59).

### Labeling SSB with Alexa Fluor 647

SSB A122C was labeled with Alexa Fluor 647 maleimide (ThermoFisher) per the manufacturer’s standard maleimide reaction protocol. Labeled protein was purified from excess fluorophore using a BioGel P-6DG desalting column followed by ion exchange chromatography using a HiTrap Q HP column (60).

### DNA annealing

Oligonucleotides were purchased from and purified by Integrated DNA Technologies (IDT) using HPLC for fluorophore-labeled oligonucleotides and PAGE for unlabeled oligonucleotides. Single-stranded DNA substrates were mixed at equal concentrations in 20 mM Tris HCl pH 7.5 and 50 mM NaCl, heated to 80°C for 5 min, and cooled to room temperature over at least 4 hours to anneal. Table S2 contains the DNA sequences of oligonucleotides that were annealed to make the substrates in Table S3.

### FRET-based helicase assay

FRET-based helicase assays were used to measure DNA duplex unwinding (11). DNA substrates (O1 and F1, Table S3) were labeled at the blunt end with a Cy3-Cy5 FRET pair. The annealed DNA (50 nM) was mixed with ATP (2 mM) in assay buffer A. Cy3 fluorescence (565 nm) was monitored continuously as a function of time while DNA, SSB (75 nM when present) and helicase were added to the cuvette sequentially. The data was analyzed using the previously published protocol (11). Three technical repeats were performed as a function of helicase concentration in Sup Fig 1 and 2.

### Gel-based helicase assay

To confirm DinG was annealing DNA in the presence of SSB ΔC1 on the overhang substrate, DNA products were measured using a gel helicase assay. Reactions contained Cy5-labeled duplex DNA (50 nM, O2), ATP (2 mM), assay buffer A, and SSB ΔC1 (75 nM). The reaction started with the addition of DinG (20 or 50 nM). Samples were removed from the reaction at 1, 5, 10, 15, 25, and 35 min and quenched with 1.5% SDS, 15 mM EDTA, and 37.5% glycerol. In the positive control (ssDNA), DinG was not added to the reaction mix and the sample was heated at 95°C for 5 min and put directly on ice. For the negative control (dsDNA), DinG was not added to the reaction mix and the sample was not denatured. Substrates were separated from products on a 10% native gel (10% acrylamide:bis solution, 19:1), which was run at 0.02 amps in a 4° C room for 20 mins. The gel was imaged with an Amersham Typhoon (Cytiva) and quantified using ImageQuantTL. Three technical repeats were performed for the gel helicase assays.

### Fluorescein quench assay

Reactions contained DNA with a fluorescein located within the duplex (50 nM, O3, O4 and F2, Table S3), ATPγS (0.5 mM), and assay buffer A. If the reaction contained wt SSB or SSB ΔC1, SSB (75 nM) was preincubated with the DNA before the addition of the helicase. A DNA-only emission spectrum from 505 to 625 nm was taken with a 495 nm excitation and 2-nm bandwidth prior to adding helicase. Then, either YoaA-χ or DinG was added to the reaction and a time-based scan was taken at 495 nm excitation and 525 nm emission. An emission spectrum was measured for the helicase-DNA complex after the signal plateaued at the same conditions as the DNA-only emission. The helicase-DNA emission spectrum was divided by the DNA-only emission spectrum at 516 nm for each trial to determine the relative quench in fluorescence. To correct for dilution, the relative helicase-DNA emission spectrum was divided by the relative DNA-only emission spectrum at 516 nm. Three technical repeats were performed for all fluorescein quench assays. As a control to show that wt SSB and SSB ΔC1 did not affect fluorescein fluorescence, the experiment was performed as above except SSB (50 nM, 75 nM, and 100 nM) was titrated onto the overhang DNA (50 nM, O4) or forked DNA (50 nM, F2) without a helicase present.

### Electrophoretic mobility shift assay (EMSA)

EMSAs were performed with dT_65_ ssDNA labeled with a Cy5 fluorophore (Table S2, S11) (50 nM), ATPγS (0.5 mM), glycerol (6.75%), and assay buffer A. Reactions containing SSB or SSB ΔC1 were pre-incubated with 75 nM SSB before the helicase was added. The helicase was titrated in concentrations ranging from 50 to 400 nM. Reactions were incubated at room temperature for 20 minutes, were loaded onto a 5% native gel (5% acrylamide:bis solution, 37.5:1) and run at 0.02 amps in a 4° C room. The gel was scanned using an Amersham Typhoon (Cytiva). EMSAs were replicated with three technical repeats.

### Single-molecule assay

Single-molecule experiments were performed on a LUMICKS C-Trap DYMO. Flow-cell passivation was performed by flowing 300 µL of the appropriate buffers in each channel, flushing each channel with 500 μL of LUMICKS passivation buffer Pluronic (diluted 1:10 in water), flushing each channel with 500 μL of LUMICKS passivation buffer BSA (diluted 1:10 in water), and then flowing 300 μL of the appropriate buffer for each channel at 1.6 bar. The microfluidic flow cell was set up with channel 1 containing 4.34 μm streptavidin-coated polystyrene beads (LUMICKS) in buffer B, channel 2 with biotinylated 20,452-nt ssDNA in buffer B (LUMICKS), channel 3 with buffer C, channel 4 with AF647-labeled SSB (20 pM), ATP (4 mM), Trolox (1 mM), GODCAT (glucose oxidase 0.54 mg/mL and catalase 0.048 mg/mL), and glucose (0.65%) in assay buffer A. If YoaA-χ or YoaA-χ R128A were used, they were also included in channel 4 at 100 nM final concentration. The overall power of the trapping laser was set to 20%. The 638 nm excitation laser was used at 1.6 μW to visualize the AF647-labeled SSB. Kymographs were collected at constant force of 10 pN with no flow, using a pixel size of 50 nm and pixel dwell time of 0.1 ms.

Kymographs were analyzed using Lakeview Pro with minimum photon counts set at 2, positional search range at 0.07 μm, maximum gap at 10 scan lines, minimum length at 10 points, expected spot size at 0.50 μm, and expected velocity at 0 μm/s. Manual refinements were performed when necessary during data analysis in Lakeview Pro to ensure track measurements were accurate. Binding events were binned into 2 second intervals. Data was exported from Lakeview Pro and the slopes of SSB movement were manually calculated.

### Structure Prediction

A model of YoaA-χ bound to ssDNA (dT_11_) was generated with AlphaFold 3 (51). The AlphaFold model of YoaA-χ-dT_11_ was superimposed on the structure of ψχ bound to a SSB peptide (PDB ID: 3SXU) to show where SSB would bind YoaA-χ (31).

### Data Availability

Data that was not shown in this article can be shared upon request.

## Acknowledgments

We thank Dr. Rafael Camerini-Otero (DinG), Dr. Michael O’Donnell (SSB), and Dr. James Keck (SSBΔC1) for providing protein expression plasmids. We also thank Elijah Newcomb and Andrea Murciano for labeling SSB with Alexa Fluor 647. We thank Alyssa Goodyear for her contribution to the fluorescein quench assays. Models were generated with BioRender. This work was supported by the National Science Foundation [MCB 1817869 to L.B.B.] and the National Institutes of Health [GM 140166 to L.B.B.] and [NIEHS F31ES034652 to S.W.P.]. The content is solely the responsibility of the authors and does not necessarily represent the official views of the National Institutes of Health.

## Author Contributions

S.J.W.P., M.J.P, and L.B.B conceived and designed the analysis. S.J.W.P., M.J.P, R.D., and K.K.H. collected the data. S.J.W.P., M.J.P, R.D., K.K.H., and L.B.B performed the analysis. S.J.W.P. and M.J.P wrote the original draft. S.J.W.P., M.J.P, R.D., K.K.H., and L.B.B reviewed and edited the draft. L.B.B supervised and acquired funding for the project.

## Ethics Declaration

The authors declare no competing interests.

## Supplementary Information

This article contains supporting information.

