## Supporting Information for "Single-stranded DNA binding protein hitches a ride with the *Escherichia coli* YoaA-χ helicase"

**Table S1:** DNA primers to create mutants

| **Primer Name** | **Sequence 5ˈ to 3ˈ** |
| --- | --- |
| χ R128A Forward Primer | ACAACTGGCGGCCGAACGCTATAAAG |
| χ R128A Reverse Primer | TTCAGAGAATCTTCATAAGGAAC |

**Table S2:** DNA oligonucleotides

| **DNA Substrate Name** | **DNA Length (nt)** | **Sequence (5ˈ to 3ˈ)** |
| --- | --- | --- |
| S1 | 85 | TTT TTT TTT TTT TTT TTT TTT TTT TTT TTT TTT TTT TTT TTT TTT TTT TTT TTT TTT TTT TTT TTA TAA AAG GGA CAT TCT GGC C-Cy3 |
| S2 | 30 | Cy5-GGC CAG AAT GTC CCT TTT ATT ACT GGT CGT |
| S3 | 20 | Cy5-GGC CAG AAT GTC CCT TTT AT |
| S4 | 30 | GGC CAG AAT GTC CCT TTT ATT ACT GG**F** CGT |
| S5 | 30 | GGC CAG AAT GTC CCT TTT ATT AC**F** GGT CGT |
| S6 | 30 | GGC CAG AAT GTC CCT TTT A**F**T ACT GGT CGT |
| S7 | 30 | GGC CAG AAT GTC CC**F** TTT ATT ACT GGT CGT |
| S8 | 30 | GGC CAG AAT G**F**C CCT TTT ATT ACT GGT CGT |
| S9 | 40 | GGC CAG AAT GTC CCT TTT ATT AC**F** GGT CGT TAC TGG TCG T |
| S10 | 95 | TTT TTT TTT TTT TTT TTT TTT TTT TTT TTT TTT TTT TTT TTT TTT TTT TTT TTT TTT TTT TTT TTA CGA CCA GTA ATA AAA GGG ACA TTC TGG CC |
| S11 | 65 | Cy5-TTT TTT TTT TTT TTT TTT TTT TTT TTT TTT TTT TTT TTT TTT TTT TTT TTT TTT TTT TTT TTT TT |
| S12 | 85 | TTT TTT TTT TTT TTT TTT TTT TTT TTT TTT TTT TTT TTT TTT TTT TTT TTT TTT TTT TTT TTT TTA TAA AAG GGA CAT TCT GGC C |

Nucleotide (nt), fluorescein (F)

**Table S3:** DNA substrates

| **Experiment** | **DNA Substrate Name** | **Oligonucleotides Annealed^1^** |
| --- | --- | --- |
| FRET-based helicase assay | O1 (overhang) | S3 and S1 |
| FRET-based helicase assay | F1 (fork) | S2 and S1 |
| Gel-based helicase assay | O2 (overhang) | S3 and S12 |
| Ruler assay | O3 (overhang) | S4, S5, S6, S7, or S8 and S10 |
| Fluorescein quench assay | O4 (overhang) | S5 and S10 |
| Fluorescein quench assay | F2 (fork) | S9 and S10 |

^1^DNA sequences for substrates are in Table S2.

Fluorescence resonance energy transfer (FRET)

**Table S4:** Quantification of relative intensity of fluorescein located 7-nt away from ds/ss junction within the duplex on a DNA substrate (50 nM) in the presence of either YoaA-χ or DinG. If wt or SSB ΔC1 (75 nM) was included in the reaction, it was preincubated on the DNA. Averages are for three experiments and error represents the standard deviation.

| **DNA Substrate** | **Helicase** | **Helicase Concentration (nM)** | **Fluorescein Relative Intensity without SSB** | **Fluorescein Relative Intensity with SSB** | **Fluorescein Relative Intensity with SSB ΔC1** |
| --- | --- | --- | --- | --- | --- |
| Overhang (O4) | YoaA-χ | 50 | 0.988 ± 0.063 | 0.919 ± 0.110 | 1.03 ± 0.021 |
| Overhang (O4) | YoaA-χ | 100 | 0.913 ± 0.106 | 0.812 ± 0.182 | 0.988 ± 0.066 |
| Overhang (O4) | YoaA-χ | 250 | 0.710 ± 0.113 | 0.609 ± 0.148 | 0.978 ± 0.019 |
| Overhang (O4) | YoaA-χ | 500 | 0.662 ± 0.154 | 0.499 ± 0.070 | 1.00 ± 0.060 |
| Overhang (O4) | YoaA-χ | 1000 | 0.457 ± 0.086 | 0.430 ± 0.029 | 0.961 ± 0.025 |
| Overhang (O4) | YoaA-χ | 1500 | 0.483 ± 0.089 | 0.370 ± 0.029 | 0.862 ± 0.038 |
| Overhang (O4) | DinG | 50 | 0.814 ± 0.096 | 0.833 ± 0.071 | 0.935 ± 0.055 |
| Overhang (O4) | DinG | 100 | 0.773 ± 0.043 | 0.664 ± 0.072 | 0.943 ± 0.069 |
| Overhang (O4) | DinG | 250 | 0.638 ± 0.088 | 0.632 ± 0.095 | 0.861 ± 0.081 |
| Overhang (O4) | DinG | 500 | 0.494 ± 0.010 | 0.572 ± 0.023 | 0.854 ± 0.046 |
| Overhang (O4) | DinG | 1000 | 0.434 ± 0.025 | 0.509 ± 0.077 | 0.753 ± 0.089 |
| Overhang (O4) | DinG | 1500 | 0.414 ± 0.073 | 0.524 ± 0.052 | 0.713 ± 0.048 |
| Fork (F2) | YoaA-χ | 50 | 1.02 ± 0.008 | 0.916 ± 0.032 | N.M. |
| Fork (F2) | YoaA-χ | 100 | 1.00 ± 0.027 | 0.918 ± 0.034 | N.M. |
| Fork (F2) | YoaA-χ | 250 | 0.939 ± 0.016 | 0.801 ± 0.046 | N.M. |
| Fork (F2) | YoaA-χ | 500 | 0.832 ± 0.027 | 0.736 ± 0.025 | N.M. |
| Fork (F2) | YoaA-χ | 1000 | 0.676 ± 0.033 | 0.565 ± 0.054 | N.M. |
| Fork (F2) | YoaA-χ | 1500 | 0.632 ± 0.016 | 0.466 ± 0.020 | N.M. |
| Fork (F2) | DinG | 50 | 0.933 ± 0.032 | 0.974 ± 0.007 | N.M. |
| Fork (F2) | DinG | 100 | 0.905 ± 0.042 | 0.910 ± 0.022 | N.M. |
| Fork (F2) | DinG | 250 | 0.766 ± 0.033 | 0.871 ± 0.102 | N.M. |
| Fork (F2) | DinG | 500 | 0.731 ± 0.039 | 0.839 ± 0.014 | N.M. |
| Fork (F2) | DinG | 1000 | 0.646 ± 0.016 | 0.754 ± 0.064 | N.M. |
| Fork (F2) | DinG | 1500 | 0.608 ± 0.037 | 0.732 ± 0.084 | N.M. |

N.M. stands for not measured.

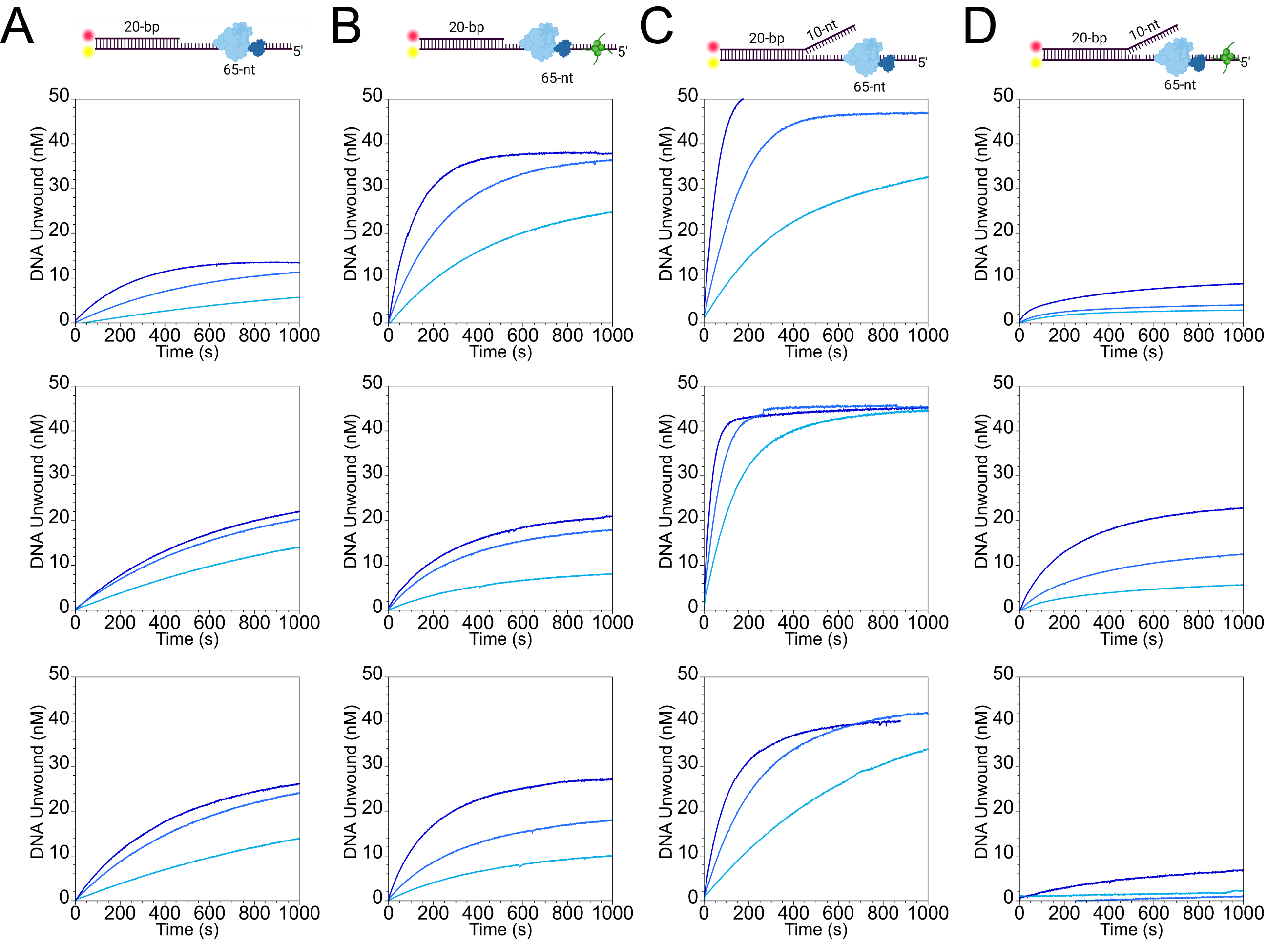

**Sup Fig 1:** Concentration range of YoaA-χ unwinding overhang and forked DNA with and without SSB. Three experiments of 5ˈ overhang DNA duplex (50 nM, O1) unwound by YoaA-χ (10 nM, 20 nM, and 50 nM) **A.** without SSB present and **B**. with SSB (75 nM) pre-bound to the DNA. Three experiments of forked DNA duplex (50 nM, F1) with a 10-nt 3ˈ overhang unwound by YoaA-χ (10 nM, 20 nM, and 50 nM) **C.** without SSB present and **D.** with SSB (75 nM) pre-bound to the DNA. Darkness in blue corresponds to increasing concentrations of YoaA-χ. On the DNA schematic, the red circle indicates Cy5, the yellow circle indicates Cy3, YoaA-χ is denoted in blue, and SSB in green.

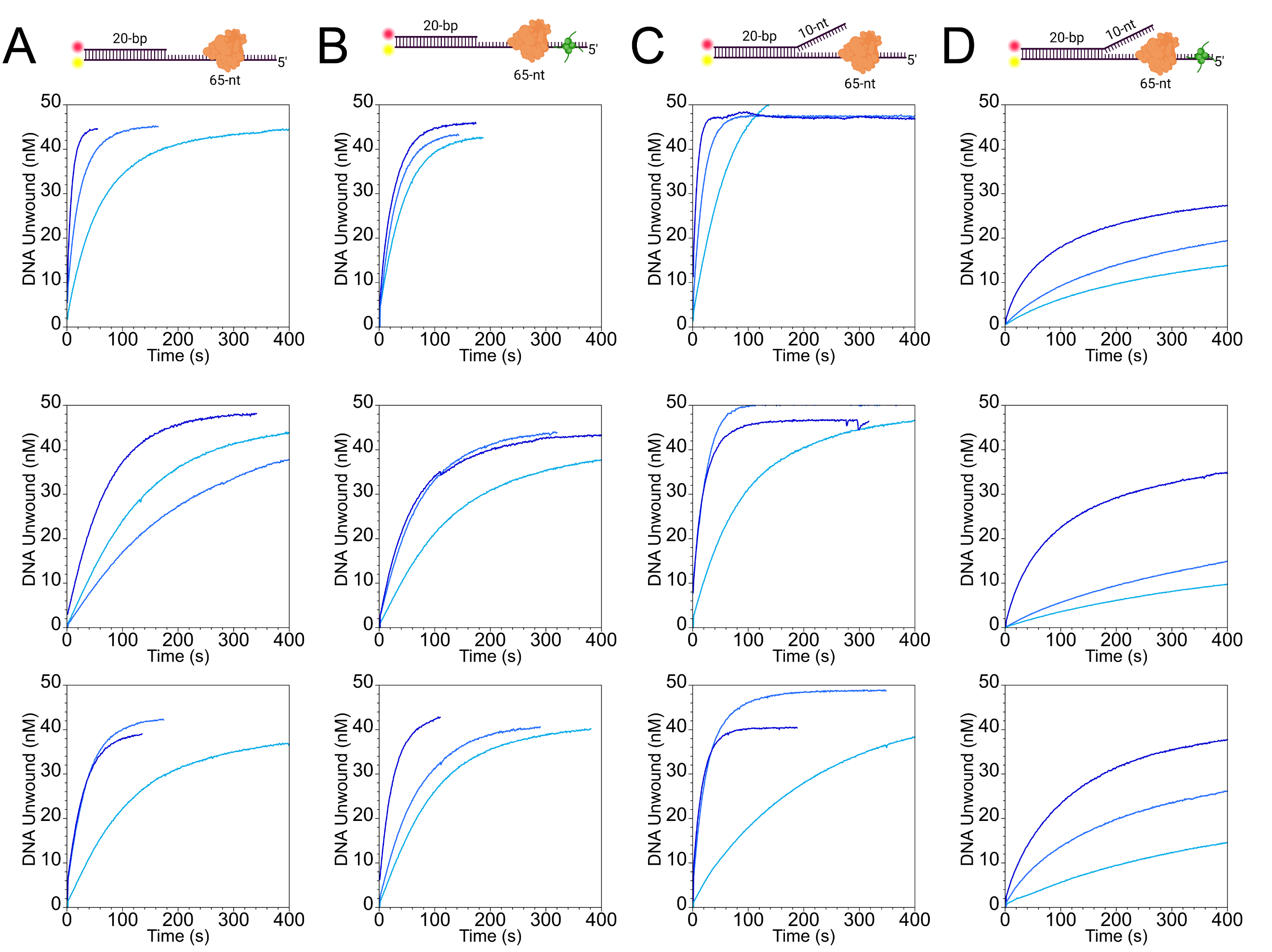

**Sup Fig 2:** Concentration range of DinG unwinding overhang and forked DNA with and without SSB. Three experiments of 5ˈ overhang DNA duplex (50 nM, O1) unwound by DinG (2 nM, 5 nM, and 10 nM) **A.** without SSB present and **B**. with SSB (75 nM) pre-bound to the DNA. Three experiments of forked DNA duplex (50 nM, F1) with a 10-nt 3ˈ overhang unwound by DinG (2 nM, 5 nM, and 10 nM)) **C.** without SSB present and **D.** with SSB (75 nM) pre-bound to the DNA. Darkness in blue corresponds to increasing concentrations of DinG. On the DNA schematic, the red circle indicates Cy5, the yellow circle indicates Cy3, DinG is denoted in orange, and SSB in green.

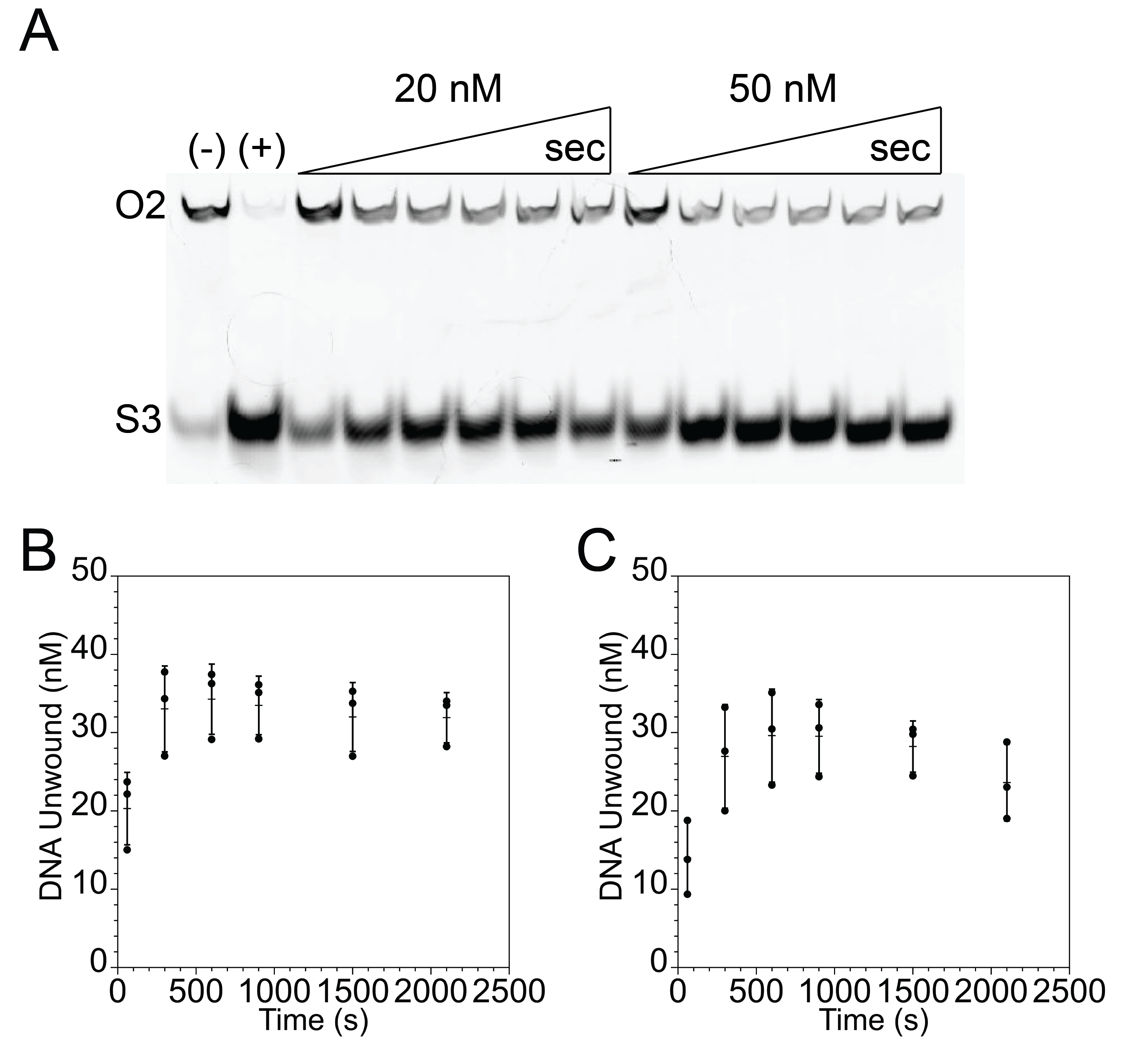

**Sup Fig 3:** Gel-based DNA helicase assay and quantification with DinG and SSB ΔC1. **A.** Representative gel of either 20 nM or 50 nM DinG added to a 5ˈoverhang DNA duplex (50 nM, O2) containing a Cy5-label on the primer prebound with 75 nM SSB ΔC1. The (-) lane contains reaction with no DinG. The (+) lane contains reaction with no DinG that was boiled at 95˚ C for 5 min and then put on ice. Time points were taken at 1, 5, 10, 15, 25, and 35 min. The experiment was performed in triplicate and a representative gel is shown. **B.** Quantification of three gel images with 50 nM DinG. **C.** Quantification of three gel images with 20 nM DinG. Dots represent individual experiments, line represents the average of three technical repeats, and error bars represent the standard deviation.

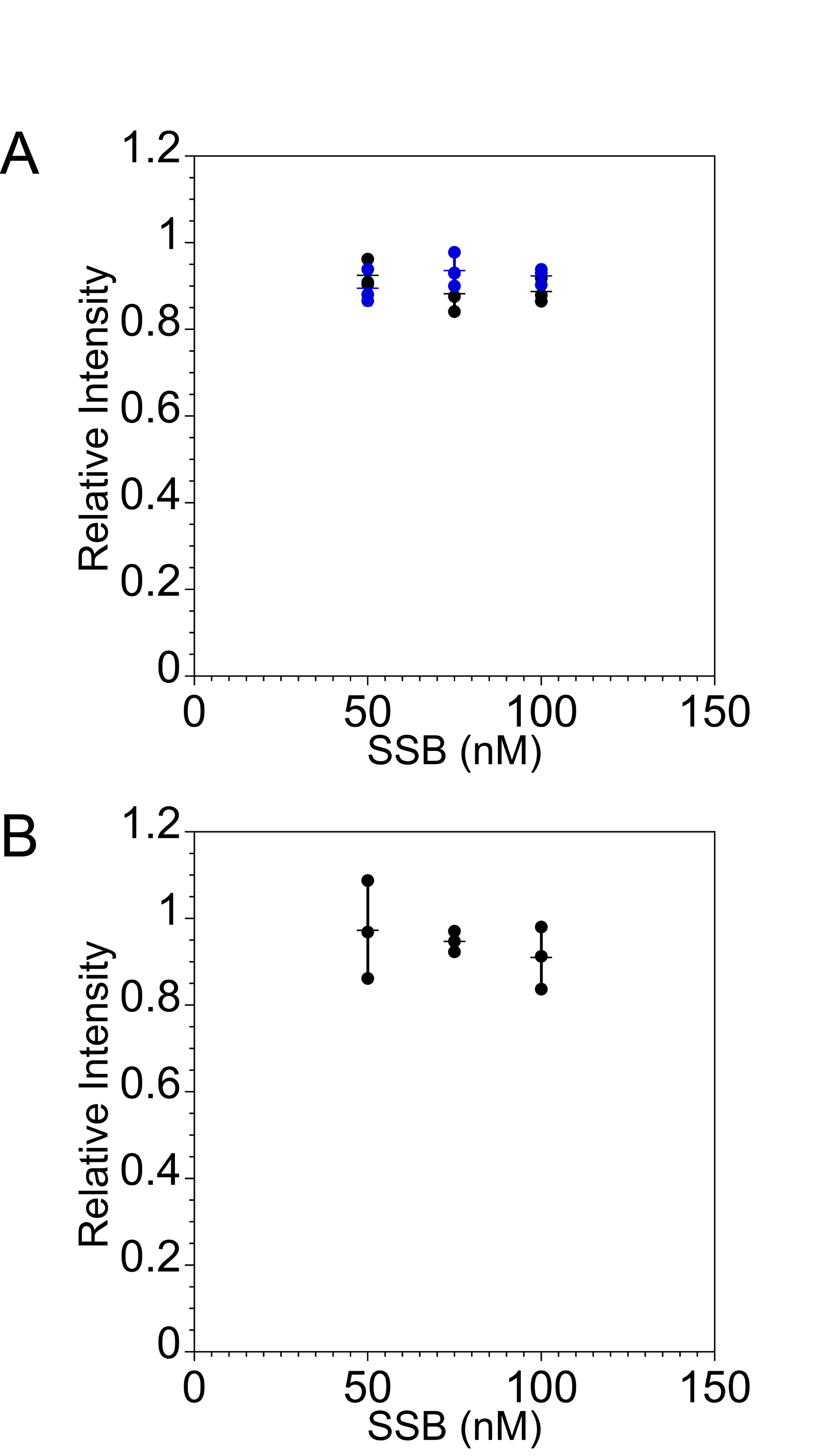

**Sup Fig 4:** SSB and SSB ΔC1 do not affect the fluorescence of a fluorescein located within the duplex 7-nt back from the ds/ss junction. **A.** Relative intensity of fluorescein located 7-nt away from ds/ss junction on overhang substrate (50 nM, O4) in the presence of either wt SSB (50 nM, 75nM, or 100 nM) (black) or SSB ΔC1 (50 nM, 75nM, or 100 nM) (blue). **B.** Relative intensity of fluorescein located 7-nt away from ds/ss junction on forked substrate (50 nM, substrate F2) in the presence of SSB (50 nM, 75nM, or 100 nM). Dots represent individual experiments, line represents the average of three technical repeats, and error bars represent the standard deviation.
